# Transcriptome remodeling and adaptive preservation of muscle protein content in hibernating black bears

**DOI:** 10.1101/2025.03.06.641932

**Authors:** Vadim B. Fedorov, Arthur Garreau, Øivind Tøien, Brian M. Barnes, Anna V. Goropashnaya

**Affiliations:** Institute of Arctic Biology, University of Alaska Fairbanks, Fairbanks, Alaska, USA

**Author notes:** **Correspondence:** Vadim B. Fedorov.

**Keywords:** bear, functional genomics, gene expression, hibernation, protein biosynthesis, RNA-seq

## Abstract

Hibernation is an energy-saving adaptation associated with physical inactivity. In contrast to most mammals, hibernating bears demonstrate limited loss of muscle mass and protein content over the prolonged periods of immobility and fasting during winter. This suggests that bears have natural adaptive mechanisms preserving muscle mass and functionality. To identify transcriptional changes that underlie molecular mechanisms attenuating muscle loss, we conducted a large-scale gene expression profiling (14,199 genes) by transcriptome sequencing in quadriceps of adult black bears, comparing hibernating animals (n=5) and summer active animals (n=5). Gene set enrichment analysis showed significant positive correlation between hibernating phenotype and expression changes of genes involved in translation, ribosome and the mTORC1 mediated signaling. In contrast, coordinated transcriptional reduction was detected for genes involved in catabolism of branched chain amino acid (BCAA) suggesting preservation of BCAA. These findings imply maintenance of protein biosynthesis through the mTORC1 signaling positively activated by availability of BCAA in muscle during hibernation. Support for this conclusion comes from overexpression of *RRAGD* and *RRAGB*, crucial regulator of the mTORC1 response to leucine availability, and up regulation of *EIF4B*, downstream target of the mTORC1 signaling. Consistent with the mTORC1 suppression of autophagy-dependent protein degradation, *MAP1LC3A* and *ULK1* were down regulated in hibernating muscle. The maintenance of protein biosynthesis and decrease in protein catabolism through the mTORC1 signaling as response to BCAA availability likely contribute to the preservation of muscle protein through prolonged periods of immobility and fasting during hibernation.

## 1. Introduction

Hibernation is an energy-saving adaptive strategy that in mammals involves suppression of metabolism and physical inactivity during periods of low food availability in seasonal environments (Carey et al. 2003). Physical inactivity decreases the mechanical load on the skeleton, which when prolonged leads to loss of muscle strength and mass in most mammalian species. Skeletal muscles are the major protein reservoir serving as a source of energy for homeostasis during periods of fasting (Sartori et al. 2021). In contrast to other mammals, hibernating species demonstrate limited loss of muscle mass and protein content over prolonged inactivity and fasting during winter (Cotton 2016). This adaptation allows hibernating mammals to maintain muscle functionality and mobility during and immediately after hibernation, thus promoting survival (Fedorov et al. 2014).

The American black bear, *Ursus americanus*, represents a well-documented example of muscle preservation throughout periods of near-complete immobility during 3-6 month hibernation season (Harlow et al., 2001). Hibernating bears maintain core body temperature above 30^◦^C but reduce whole – body metabolic rate by 75% (Tøien et al. 2011). They do not eat, drink, defecate nor urinate (Nelson et al. 1973). Although a moderate (10%) decrease in muscle protein content reported for lactating female bears (Tinker et al. 1998), no loss of muscle mass and protein was detected over winter in hind limb muscles of hibernating American black bears (Lundberg et al. 1976; Koebel et al. 1991; Harlow et al. 2001; Lohuis et al. 2007a, b). Similarly, Asiatic black bears, *U. thibetanus*, and brown bears, *U. arctos*, maintain muscle mass and fiber composition during hibernation (Miyazaki et al. 2022; Hershey et al. 2008). This suggests that several species of bears have evolved natural adaptive mechanisms preserving muscle mass and functionality over prolonged periods mechanical unloading and fasting. The molecular mechanisms underlying this protective musculoskeletal adaptation remain poorly understood.

Previous studies of transcriptional changes on the sub-genomic scale in skeletal muscle of hibernating black bears revealed an elevated expression of genes involved in protein biosynthesis and ribosome biogenesis, but detected no directional changes for genes of protein catabolism (Fedorov et al. 2009; 2014). These findings are in a contrast to the transcriptional suppression of protein biosynthesis genes and elevated expression of catabolic genes during disuse that are the hallmarks of muscle atrophy in non-hibernating mammals (Lecker et al. 2004; Abadi et al. 2009). These results led to the working hypothesis that transcriptional induction of protein biosynthesis genes contributes to molecular mechanisms maintaining muscle mass and strength during physical inactivity of hibernation (Fedorov et al. 2014). However, the limited genome coverage obtained with custom cDNA microarray approaches used previously in black bears reduced the power of pathway enrichment analysis.

Support for the transcriptional induction of protein biosynthesis in hibernating muscle comes from the recent genome-wide transcriptome sequencing study in brown bears showing significant positive correlation between expression values of anabolic genes and hibernating phenotype (Jansen et al. 2019). In addition, this study revealed coordinated transcriptional suppression of urea cycle genes in liver of hibernating brown bears that suggests redirection of amino acids from catabolic pathways to active protein biosynthesis.

In the present study, to increase genome coverage we conducted unbiased transcriptome sequencing (RNA-seq) to quantify gene expression changes in the muscle of hibernating black bears in comparison to summer active bears. We used genome-wide expression data to conduct gene set enrichment analysis identifying functional groups of co-regulated genes and reveal the biological significance of transcriptional changes for muscle preservation. We also selected from our data set genes with known functional relation to muscle homeostasis in non-hibernating mammals and considered transcriptional changes in light of muscle maintenance during the disuse and fasting of hibernation. Finally, to reveal general transcriptional program in muscle of hibernating bears, we conducted pathway enrichment analysis of differentially expressed genes that are common in black and brown bears (Jansen et al. 2019).

## 2. Materials and Methods

### 2.1 Animals

All protocols for animal work were approved by the University of Alaska Fairbanks, Institutional Animal Care and Use Committee (IACUC nos. 02-39, 02-44, 05-55, and 05-56). Black bears (51.2-226 kg) were captured in Alaska by Alaska Department of Fish and Game during May– July, transferred to Fairbanks and held individually in an outdoor facility. Only males > 2 years old were selected in these experiments to minimize effects of sex and age of intra-group variation in gene expression. Summer active bears (n = 5) were euthanized after fasting for 24 hours and sampled for tissues between June and early October. Hibernating bears (n = 5) were without water and food since October 27 and were sacrificed for tissue collection between March 1-27, about one month before their expected emergence from hibernation. Monitoring of physiological conditions of hibernating bears was described in Fedorov et al. (2009) and Tøien et al. (2011, 2015). Briefly, core temperature, EKG and EMG were recorded with radio telemetry. In late November, the bears were moved into closed individual dens with straw nest material where air was drawn through the dens, and, oxygen consumption and respiratory quotient were evaluated with an open flow respirometry system (Tøien 2013). Bears were euthanized by an intravenous injection of pentobarbital, and was death assessed by termination of heart beats as detected with a stethoscope. Tissues including the hind limb muscle (quadriceps femoris) were sampled immediately and frozen in liquid nitrogen within 12 min of death.

### 2.2 RNA preparation and sequencing

Frozen skeletal muscles (approximately 30 mg) were homogenized in 2 ml Lysing Matrix S tubes with beads (MP Biomedical) containing 300µL of RLT buffer (Qiagen) using a Mini-Beadbeater-1 (BioSpecProducts) for 100 sec at 2500 oscillations/minute. Total RNA was isolated with a RNeasy Fibrous mini kit (Qiagen) according to manufacturer’s instructions. Any remaining genomic DNA was removed with a DNase I (Qiagen) treatment. The RNA quality and concentration were obtained with a TapeStation 4200 (Agilent Technologies) and a Nanodrop ND-1000. The average RIN value was 8.0, with no differences between sample groups. Ribosomal RNA content was calculated as a ratio of amount of 18S and 28S ribosomal RNA over total RNA. Total RNA was used for strand specific mRNA sequencing on DNBSEQ platform PE150 (BGI Americas Corporation).

### 2.3 Data analysis

On average, 70 million 150 bp paired-end reads per sample were obtained and mapped to the reference genome of *U. americanus* (black bear, GCF_020975775.1) using CLC Genomics Workbench software (v.20.0.4, https://www.qiagenbioinformatics.com). In total, 96.01% of reads were mapped in pairs with the following options: mismatch cost = 2, insertion cost = 3, deletion cost = 3, length fraction = 0.9, deletion fraction = 0.9. Genes expression values were determined using total counts of reads mapped in pair to the exons and normalized for library size using the TMM method (Robinson and Oshlack 2010). Only genes with at least 2 reads across all samples were included in further analysis. The dispersion parameter of normalized read counts for each gene was estimated with the multi-factorial EdgeR method implementing negative binomial Generalized Linear Model (Robinson et al. 2010). The Wald test was applied to assess gene expression differences between the hibernating and summer active bears. For each gene, the false discovery rate (FDR) was determined using the procedure described by Benjamini and Hochberg (1995). Genes were considered differentially expressed if FDR was 0.05 or less and gene expression fold change was greater than 1.5. Gene set enrichment analysis (GSEA) was applied to estimate enrichment in gene sets corresponding to biological processes or metabolic pathways (Subramanian et al. 2005). The software uses, as an input, the whole list of genes pre-ranked according to their fold change obtained in pairwise comparison between hibernation and summer active bears. GSEA assessed overrepresentation of up and down regulated genes in gene sets and calculated enrichment score (ES) to estimate the degree to which a gene was overrepresented at the extremes (up regulated at the top and down regulated at the bottom) of the entire ranked list of genes. ES was then normalized to adjust for the size of the genes sets, thus providing normalized ES (NES). High positive values of NES correspond to a large number of up regulated genes in a gene set, low negative NES values reflect a large number of down regulated genes, accordingly. A premutation test based on gene set was used to estimate the statistical significance of the NES. A gene set was considered up or down regulated with |NES| > 1.5 and FDR < 0.05. Gene sets corresponding to biological function or metabolic pathways were obtained from Molecular Signatures Database (http://www.broadinstitute.org/gsea/msigdb/index.jsp) that included Reactome, KEGG and WikiPathways gene set collections.

To reveal general transcriptional programs in muscle of hibernating bears, we conducted pathway enrichment analysis of differentially expressed genes that are common in black and brown bears (Jansen et al. 2019) by using Enrichr (Chen et al.2013; https://maayanlab.cloud/Enrichr/#). Lists of significant differentially expressed genes common for both species (Table 1S) were input to Enrichr to estimate, with the Fisher exact test, significance of elevated proportion of up or down regulated genes involved in specific biological function or pathway as compared to random expectation.

**Table 1.**
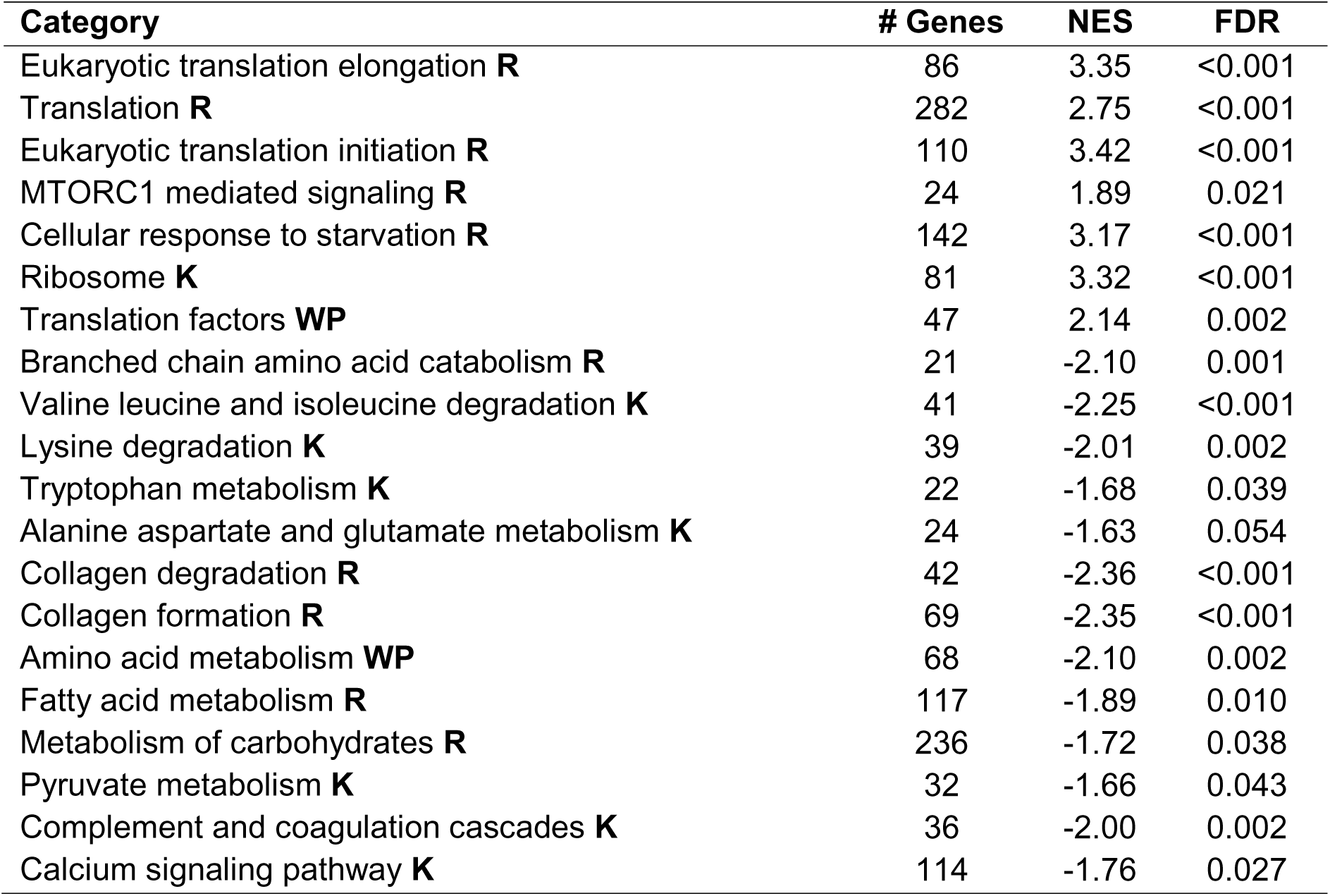
Gene set enrichment (GSEA) for selected Reactome (R), KEGG (K) and WikiPathways (WP) gene sets. Positive NES (normalized enrichment score) values indicate positive correlation between gene expression values and hibernating phenotype, negative scores indicate negative correlation. FDR is the false discovery rate.

| Category | # Genes | NES | FDR |
| --- | --- | --- | --- |
| Eukaryotic translation elongation **R** | 86 | 3.35 | <0.001 |
| Translation **R** | 282 | 2.75 | <0.001 |
| Eukaryotic translation initiation **R** | 110 | 3.42 | <0.001 |
| MTORC1 mediated signaling **R** | 24 | 1.89 | 0.021 |
| Cellular response to starvation **R** | 142 | 3.17 | <0.001 |
| Ribosome **K** | 81 | 3.32 | <0.001 |
| Translation factors **WP** | 47 | 2.14 | 0.002 |
| Branched chain amino acid catabolism **R** | 21 | -2.10 | 0.001 |
| Valine leucine and isoleucine degradation **K** | 41 | -2.25 | <0.001 |
| Lysine degradation **K** | 39 | -2.01 | 0.002 |
| Tryptophan metabolism **K** | 22 | -1.68 | 0.039 |
| Alanine aspartate and glutamate metabolism **K** | 24 | -1.63 | 0.054 |
| Collagen degradation **R** | 42 | -2.36 | <0.001 |
| Collagen formation **R** | 69 | -2.35 | <0.001 |
| Amino acid metabolism **WP** | 68 | -2.10 | 0.002 |
| Fatty acid metabolism **R** | 117 | -1.89 | 0.010 |
| Metabolism of carbohydrates **R** | 236 | -1.72 | 0.038 |
| Pyruvate metabolism **K** | 32 | -1.66 | 0.043 |
| Complement and coagulation cascades **K** | 36 | -2.00 | 0.002 |
| Calcium signaling pathway **K** | 114 | -1.76 | 0.027 |

## 3. Results

Out of 70 million paired-end 150 base pair sequencing reads generated for each sample, 88% reads were mapped in pairs to exons of the black bear reference genome. Transcriptome sequencing detected expression of 14,199 genes with at least two paired-end reads mapped to exons in each sample; that is about 56% of annotated genes in the reference genome. Total 1,013 genes were differentially expressed (FDR<0.05) between hibernating and summer active bears. Most, 677 genes, were down regulated and 336 genes were over expressed in hibernating muscle (Table 1S).

### 3.1 Gene set enrichment analysis in hibernating black bears

Gene set enrichment analysis (GSEA) identifies functional groups of co-regulated genes by estimating correlation and its significance between expression changes of genes involved in biological process or pathway and hibernating phenotype as compared to summer control. To estimate enrichment, GSEA accounts for all of the genes with expression detected in the experiment, not only those above arbitrary cutoffs for significance of expression differences, false discovery rate and fold-change (Subramanian et al. 2005).

GSEA with Reactome database showed significantly elevated proportion of overexpressed genes involved in translation (protein biosynthesis) that includes eukaryotic translation initiation and elongation (Table 1) in muscle of hibernating bears. Ribosomal proteins and translation factors are the main groups of genes involved in translation. Taken separately, expression changes of ribosome proteins (KEGG) and eukaryotic translation factors (WikiPathways) also showed significant positive correlation to the hibernating phenotype (Table 1). An important regulator of protein biosynthesis, the mTORC1 mediated signaling pathway also demonstrated significant enrichment by up regulated genes. Notably, the set of genes known to be down regulated in cellular response to starvation in non-hibernating mammals was enriched by up regulated genes during fasting of hibernation.

During hibernation, genes involved in a number of metabolic gene sets demonstrated transcriptional suppression (Table 1). An important finding in light of muscle protein turnover is that genes involved in branched chain amino acids (BCAA) catabolism, and more extensive KEGG gene set of valine, leucine and isoleucine degradation, as well as lysine degradation showed transcriptional decrease in hibernating muscle. Similarly, down regulated genes were over represented in the gene sets of amino acid metabolism, tryptophan metabolism, and alanine, aspartate, glutamate metabolism (Table 1). In addition to suppression of amino acid metabolism, we detected significant enrichment by down regulated genes for metabolism of fatty acid, carbohydrates, pyruvate, collagen as well as complement-coagulation cascades and calcium signaling pathway.

### 3.2 Enrichment analysis of differentially expressed genes shared between muscles of hibernating black and brown bears

We used a comparative approach and conducted pathway enrichment analysis of differentially expressed genes shared between black and brown bears (Jansen et al. 2019) to reveal common transcriptional program of muscle homeostasis during hibernation. Out of 276 differentially expressed genes shared between the two species, 128 genes were up regulated and 148 genes down regulated (Table 1S) in both. Lists of differentially expressed genes common for both species were used for Enrichr analysis.

Overall, results of comparative enrichment analysis in two species of bears are consistent to GSEA finding in hibernating black bears. The proportion of over expressed genes is significantly elevated among genes involved in translation for both species during hibernation (Table 2). Only one member of the mTORC1 mediated signaling, *EIF4B*, was shared between species and this gene was up regulated in both species (Table1S). Down regulated genes were over represented in the categories of BCAA catabolism, lysine degradation and metabolism of amino acids (Table 2). A suite of gene sets involved in metabolism, fatty acid betta oxidation, collagen formation demonstrated significant transcriptional suppression in both species during hibernation.

**Table 2.** Comparative gene set enrichment analysis (Enrichr) of differentially expressed genes shared between black and brown bears (Jansen et al. 2019) with Reactome (R) and KEGG (K) collections. Overlap – number of detected differentially expressed genes out of total gene number in gene set. P-value of the Fisher exact test adjusted for multiple tests.

| Category | Overlap | Adjusted P-value | Differentially Expressed Genes |
| --- | --- | --- | --- |
| *Up regulated genes* |  |  |  |
| Translation **R** | 14/281 | 3.35E-07 | *RPL30; SSR4; RPL12; MRPL23; RPS25; TRMT112; EIF3G; EIF3H; RPL38; RPL22L1; UBA52; RPS21; EIF4B; RPS23* |
| Eukaryotic Translation Termination **R** | 11/90 | 5.08E-07 | *TRMT112; RPS25; RPL30; RPL12; RPL38; RPL22L1; UBA52; RPS21; RPS23* |
| Eukaryotic Translation Elongation **R** | 8/90 | 4.31E-06 | *RPS25; RPL30; RPL12; RPL38; RPL22L1; UBA52; RPS21; RPS23* |
| Translation Initiation Complex Formation **R** | 6/57 | 4.28E-05 | *RPS25; EIF3G; EIF3H; RPS21; EIF4B; RPS23* |
| Cellular Response To Starvation **R** | 8/153 | 1.39E-04 | *RPS25; RPL30; RPL12; RPL38; RPL22L1; UBA52; RPS21; RPS23* |
| Ribosome **K** | 9/158 | 9.01E-05 | *RPS25; RPL30; RPL12; RPL38; RPL22L1; MRPL23; UBA52; RPS21; RPS23* |
| **Down regulated genes** |  |  |  |
| Branched-chain Amino Acid Catabolism **R** | 6/21 | 1.47E-06 | *ACAD8; MCCC2; BCKDHB; MCCC1; DBT; BCAT2* |
| Valine, leucine and isoleucine degradation **K** | 11/48 | 1.06E-08 | *ACAD8; MCCC2; OXCT1; DBT; BCKDHB; MCCC1; PCCB; ABAT; BCAT* |
| Metabolism Of Amino Acids And Derivatives **R** | 14/364 | 3.17E-05 | *MCCC2; MPST; ACAD8; MCCC1; BCKDHB; GADL1; GPT; DHTKD1; ALDH4A1; PXMP2; DBT; GLUL; AASS; BCAT2* |
| Lysine degradation **K** | 4/63 | 1.58E-02 | *COLGALT2; PLOD1; AASS; DHTKD1* |
| Alanine, aspartate and glutamate metabolism **K** | 4/37 | 3.48E-03 | *ALDH4A1; ABAT; GPT; GLUL* |
| Metabolism **R** | 46/2049 | 9.83E-10 | *ALDH1L1; GADL1; FITM1; ACSM5; GPT; ENO1; SAMHD1; DHTKD1; GYS1; ACADL; PHKG1; OXCT1; DBT; ARSI; GPAT3; SLC38A3; SLC37A4; ACSS1; GLUL; CA14; AASS; MCCC2; MPST; ACAD8; DGAT2; ELOVL5; EPHX2; ACOT11; MCCC1; BCKDHB; ACSL6; DHCR24; GPCPD1; PTGR1; HSPG2; SUMF2; ALDH4A1; MLXIPL; PHOSPHO1; GCLC; FASN; PXMP2; PCCB; PLBD1; HAGH; BCAT2* |
| Fatty Acid Metabolism **R** | 8/173 | 1.38E-03 | *ACADL; ELOVL5; EPHX2; FASN; ACOT11; PCCB; ACSL6; PTGR1* |
| Mitochondrial Fatty Acid Beta-Oxidation **R** | 3/36 | 3.27E-02 | *ACADL; ACOT11; PCCB* |
| Pyruvate metabolism **K** | 4/47 | 7.77E-03 | *PC; ME2; HAGH; ACSS1* |
| Collagen Formation **R** | 8/90 | 2.76E-05 | *COLGALT2; COL15A1; COL4A2; COL4A1; P4HA2; COL4A4; COL4A3; PLOD1* |

### 3.3 Transcriptional changes of selected genes involved in muscle homeostasis in the black bear

In addition to the gene set enrichment analysis, we also reviewed significant expression differences (FDR<0.05; Table 1S) or lack of changes for individual genes known to be important for muscle protein metabolism in mammals. The activated mTORC1 mediated signaling has downstream target involved in protein biosynthesis, translation initiation factor *EIF4B* (FC=1.95) that was up regulated in hibernating muscle. The upstream master stimulator of the mTORC1 signaling, insulin growth factor 1, *IGF1* was under expressed during hibernation (FC=-2.94). In contrast, the crucial activators of the mTORC1 response to leucine availability (Takahara et al. 2020), Ras-related GTP-binding protein D, *RRAGD*, was significantly over expressed (FC=2.12) and *RRAGB* demonstrated tendency for up regulation (FC=1.65; FDR=0.098).

Among BCAA catabolism genes demonstrating coordinated transcriptional reduction during hibernation, branched chain aminotransferase, *BCAT2* (FC=-1.67), catalyzes the first transamination reaction of BCAA, branched chain keto acid dehydrogenase, *BCKDHB* (FC=-2.37) is involved in the second rate-limiting irreversible step in BCAA catabolism and *MCCC1* (FC=-2.10), *MCCC2* (FC=-1.87) catabolize specifically leucine (Mann et al. 2021).

Similar to metabolism of amino acids, genes involved in fatty acid, carbohydrate metabolisms and energy production showed transcriptional suppression during hibernation. Under expression of fatty acid synthase, *FASN* (FC= −7.40), acetyl-CoA carboxylase, *ACACA* (also known as *ACC*; FC=-3.93), fatty acid elongases 5 and 6, *ELOVL5* (FC=-3.22) and *ELOVL6* (FC=-5.89), and acyl-CoA dehydrogenase, *ACADL* (FC=-2.24) implies down regulation of both fatty acid biosynthesis and catabolism (mitochondrial beta-oxidation). Transcriptional reduction of muscle specific hexokinase 2, *HK2* (FC=-1.67), enolase, *ENO1* (FC=-2.23), pyruvate carboxylase, *PC* (FC=-1.71) and glucose transporter *SLC2A1* (FC=-5.88) suggests decrease in glycolysis in hibernating muscle.

During hibernation, we detected transcriptional reduction or no expression changes for key genes involved in protein degradation through autophagy and ubiquitin proteolysis. Among autophagy related genes, microtubule associated protein 1 light chain, *MAP1LC3A* (also known as *LC3A*, FC=-2.46) and unc-51 like autophagy activating kinase, *ULK1* (FC=-1.69), were both down regulated while *ATG13*, *BECN1* showed no changes in expression.

OUT deubiquitinase 1, *OTUD1*, was the most over expressed gene (FC=12.37) in hibernating muscle. This enzyme cleaves ubiquitin linkages, negating the action of ubiquitin ligases in proteasome protein degradation and suppressing apoptosis (Oikawa et al., 2022). Two ubiquitin ligases, muscle atrophy markers and key members of proteasome degradation, *FBXO32* (also known as *Atrogin-1*, *MAFBX*) and *TRIM63* (also known as *MURF1*) as well as their upstream activators, *FOXO1* and *FOXO3* did not show differences in expression.

### 3.4 RNA content

Total RNA concentration and ribosomal RNA content were measured to assess the translational capacity of bear skeletal muscles. Total RNA amount in muscles was not different in hibernating black bears compared to summer active (Fig. 1A). Ribosomal RNA content was also not different during hibernation compared to summer active bears (Fig. 1B), revealing no decrease in translational capacity of muscles during hibernation.

**Figure 1.**
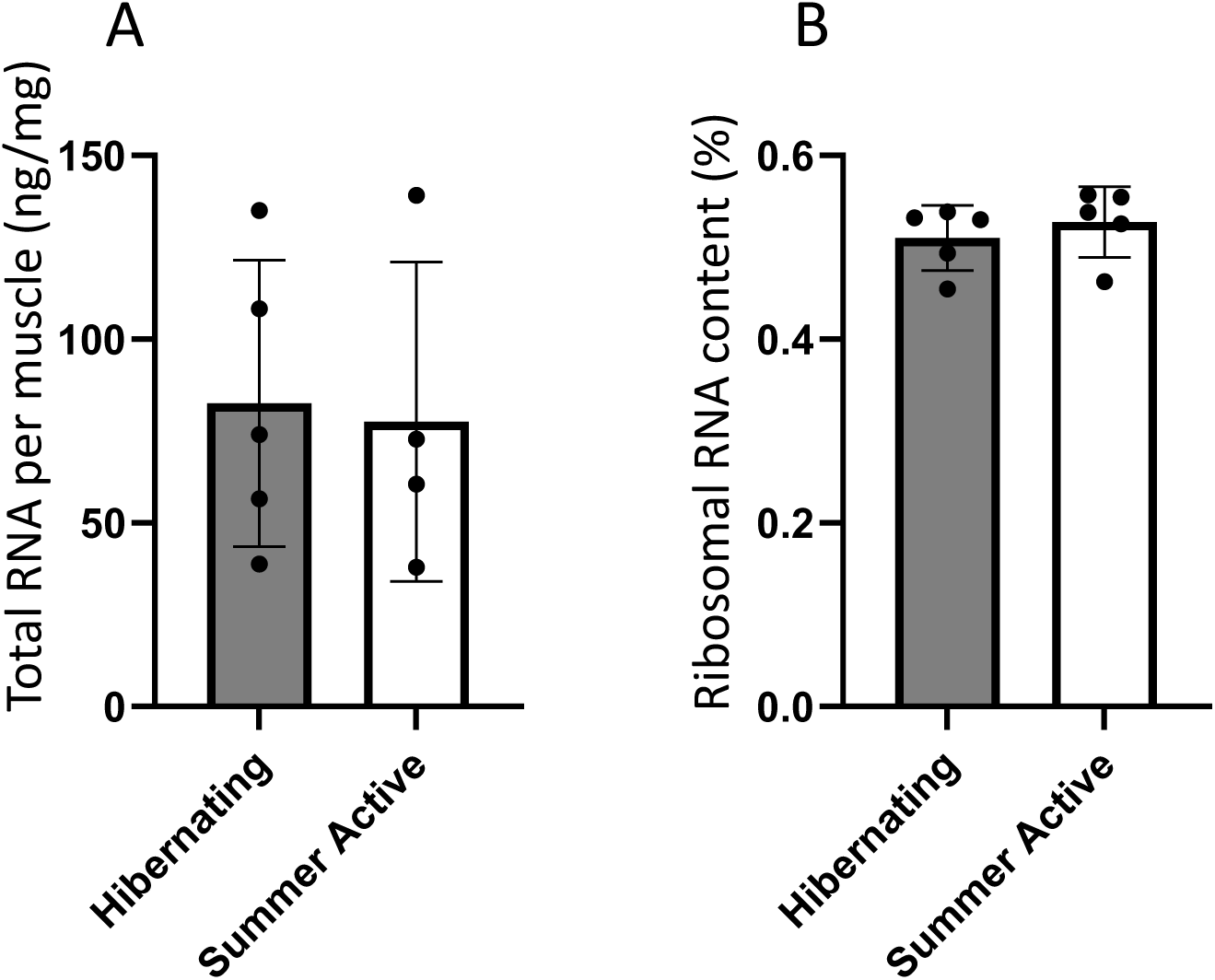
Total RNA concentration per muscle mass (A) and ribosomal RNA ratio in total RNA content inferred from 18S and 28S amount (B) in muscle of black bears during hibernation and summer seasons.

## 4. Discussion

This study reveals coordinated transcriptional changes comparing hibernating with summer active black bears that include induction of protein biosynthesis genes and suppression of genes involved in metabolism in muscle. These changes likely contribute to maintenance of muscle protein content and preservation of muscle mass during prolonged inactivity, fasting, and metabolic suppression in hibernating bears. Preservation of protein content is supported by observations that average respiratory quotient of hibernating black bears remained very constant at 0.72 in mid hibernation (Tøien 2013, Tøien et al. 2015), indicating almost exclusively use of fat as energy source with no protein breakdown.

In humans, disuse induced muscle loss results primarily from decrease in protein biosynthesis (Nunes et al. 2022) that is reflected in transcriptional down regulation of genes involved in translation (Abadi et al. 2009). In contrast to non-hibernating mammals, transcriptional up regulation of protein biosynthesis genes reported here for bears implies active anabolism leading to the prevention of muscle loss during immobility in hibernation. Our results also revealed coordinated under expression of genes involved in metabolism and degradation of amino acids during hibernation. In line with these findings, transcriptional down regulation was previously reported for the urea cycle genes in the liver of both hibernating black bears (Fedorov et al. 2009) and brown bears (Jansen et al. 2019). Reduction in urea production is supported by the decrease in the urea concentration in blood of hibernating bears attributed to both low production and urea recycling through the gut microbiome (Barboza et al. 1997; Hissa et al. 1998; Nelson 1980). Under fasting during hibernation with no dietary intake of proteins, reduced catabolism of amino acids and urea production suggest redirection of amino acids from catabolism and amino group utilization through the urea cycle to active protein biosynthesis.

Ribosomal proteins are major group of protein biosynthesis gene sets and these gene products are structural constitutes of ribosomes involved in ribosome biogenesis. The transcriptional induction of a number of ribosomal proteins reported here implies maintenance of ribosome content that contributes to overall translational capacity (Chaillou et al. 2014) in hibernating muscle. Consistent with maintenance of ribosomes, we found no decrease in total RNA concentration and stable content of ribosomal RNA in muscle of hibernating compared to summer active black bears. Similarly, no loss of total RNA was reported specifically in muscle of hibernating brown bears (Jansen et al. 2019).

The mTORC1 mediated signaling promotes protein biosynthesis and suppresses protein degradation through the inhibition of autophagy (Takahara et al. 2020). Although protein phosphorylation events are the major factors regulating the mTORC1 signaling (Kim and Guan 2019), transcriptional induction of this pathway detected here suggests activation of the mTORC1 signaling that positively regulates protein biosynthesis in muscle of hibernating bears. Insulin growth factor (*IGF1*) is an anabolic factor that activates the mTORC1 through PI3K-dependent activation of *AKT* (Takahara et al. 2020). Our results do not suggest activation the mTORC1 through the IGF1-PI3K-AKT axis during hibernation, as *IGF1* was down regulated and no expression changes were detected for *PI3K* and *AKT*.

Apart from activation by insulin growth factor, the mTORC1 signaling is stimulated by availability of BCAA (Takahara et al. 2020). Coordinated transcriptional suppression of BCAA catabolism genes suggests preservation of BCAA in muscle of hibernating bears. Taken together, our findings imply activation of protein biosynthesis through the mTORC1 signaling positively stimulated by availability of leucine in muscle during hibernation (Fig. 2). Support for this inference comes from overexpression of Ras-related GTP-binding protein B and D genes (*RRAGB*; *RRAGD*), crucial activators of the mTORC1 in response to leucine availability and up regulation of eukaryotic translation initiation factor 4B gene (*EIF4B*), downstream target of the mTORC1 that initiates protein biosynthesis.

**Figure 2.**
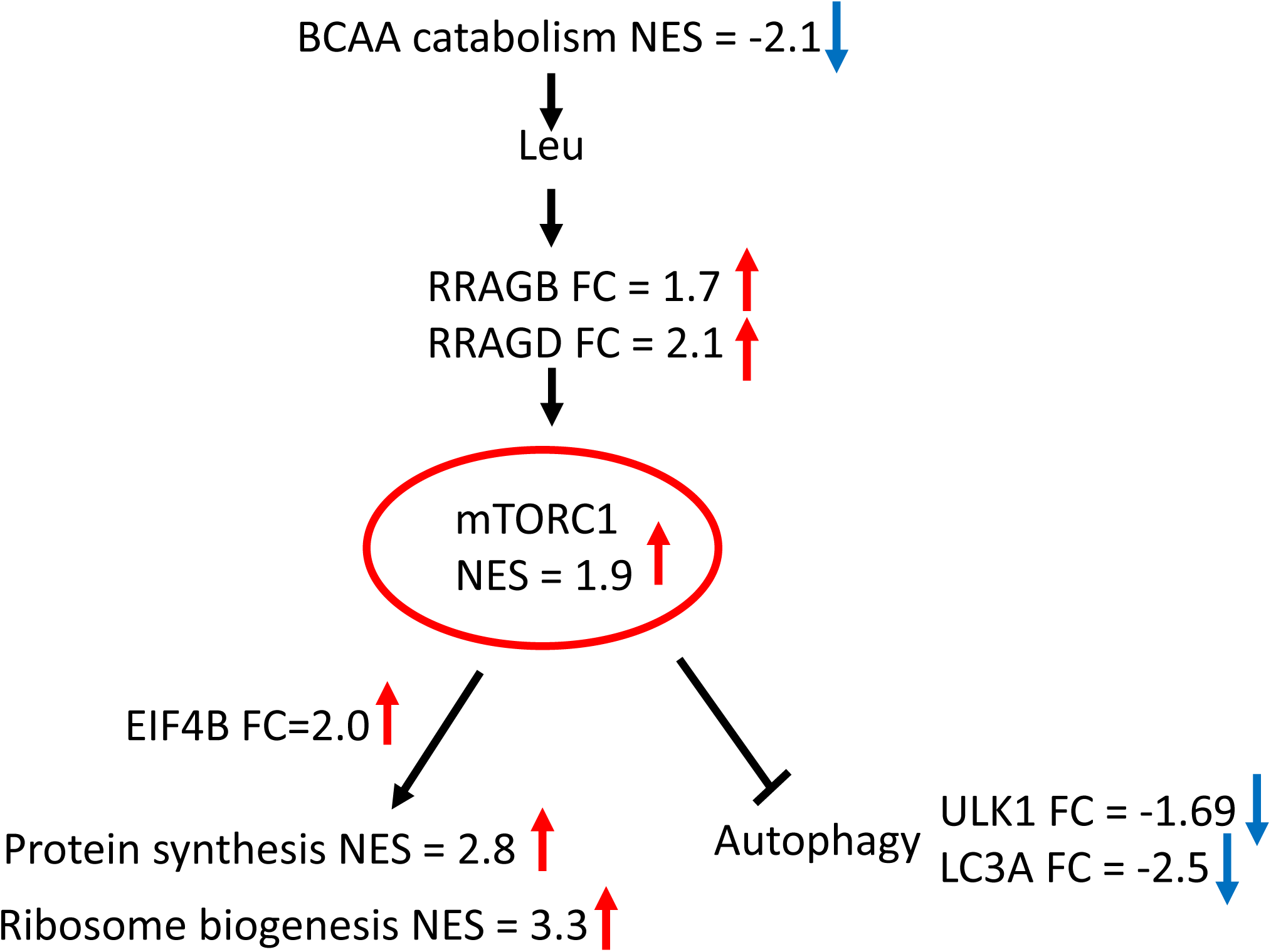
Activation of the mTORC1 mediated signaling by BCAA availability enhances protein biosynthesis and suppress autophagy (Takahara et al. 2020) in muscle of hibernating black bears. FC—fold change in expression of gene in muscle of hibernating black bears, NES—GSEA normalized enrichment score for gene set (Table 1), red arrow—up-regulation, blue arrow—down-regulation.

Gene set of the cellular response to starvation includes RAS-related GTPases, sensors of BCAA availability, and members of the mTORC1 signaling as well as a number of ribosomal proteins involved in protein biosynthesis. These genes demonstrated coordinated down regulation in response to BCAA deficiency resulting from starvation in non-hibernating mammals (Kim and Guan 2019; Liu and Sabatini 2020). Conversely, transcriptional induction of genes involved in response to starvation reported here (Tables 1, 2) for the two species of bears indicates reverse molecular effect of nutrient deficiency during hibernation. Transcriptional suppression of BCAA catabolism is likely a key factor preserving BCAA content and, thus, maintaining protein biosynthesis through supporting the mTORC1 signaling during fasting in hibernation.

In addition to promotion of protein biosynthesis, activation of the mTORC1 signaling suppresses autophagy-dependent protein degradation. Consistent with reduction of autophagy, direct downstream target of the mTORC1 (Sartori, Romanello, and Sandri 2021), unc-51 like autophagy activating kinase gene (*ULK1*), and microtubule associated protein 1 light chain gene (*MAP1LC3A*) were both down regulated in hibernating muscle. This finding is in contrast to transcriptional up regulation of *ULK1* and induction of autophagy as response to fasting in muscle of non-hibernating mammals (Galves et al. 2023).

A recent study of expression changes at protein level for selected genes involved in the mTORC1 signaling does not exclude its importance for maintaining protein biosynthesis in muscle of hibernating Asiatic black bears (Miyazaki et al. 2022). Although phosphorylation ratio of ribosomal protein *S6* (*RPS6*), downstream effector of the mTORC1, was decreased, the total quantity of *RPS6* protein was significantly elevated in muscle during hibernation. Similar to our finding, transcriptional reduction of *MAP1LC3A*, the key gene in the autophagy-lysosome pathway, was observed in muscle of hibernating Asiatic black bears (Miyazaki et al. 2022) and brown bears (Jansen et al. 2019). Suppression of autophagy dependent protein degradation likely contributes to muscle protein preservation in three species of bears during immobility and fasting in winter hibernation.

Besides autophagy, the ubiquitin proteolysis is a major pathway of muscle protein degradation during disuse in non-hibernating mammals (Taillandier et al. 1996; Solomon and Goldberg 1996). Our study did not detect expression changes of *FBXO32* and *TRIM63* ubiquitin ligases, key components in the proteasome protein degradation and biomarkers of muscle atrophy (Bodine et al. 2001). This finding implies no increase in muscle protein degradation during winter hibernation. Stable expression of these atrogenes in muscle of hibernating brown bears (Jansen et al. 2019) and their transcriptional suppression in hibernating Asiatic black bears (Miyazaki et al. 2022) were considered as factors preventing pro-atrophic protein degradation and, thus, preserving muscle mass during hibernation.

Similar to suppression of amino acid metabolism, we observed down regulation or no expression changes for genes involved in metabolism of carbohydrates and fatty acids during hibernation. This finding is consistent with the energy saving due to suppression of whole-body metabolism (Tøien et al. 2011, 2015) and transcriptional reduction of genes involved in fuel metabolism reported in muscle of hibernating black, brown and Asiatic bears (Fedorov et al. 2014; Jansen et al. 2019; Miyazaki et al. 2022). Gene expression changes at protein level suggest decrease in fatty acid oxidation but maintenance of glycolysis in muscle of hibernating brown bears (Chazarin et al., 2019). However, transcriptional down regulation of key glycolytic genes (*ENO1*, *HK2*, *PC*) observed here implies decline in glycose utilization in muscle of hibernating black bears.

Hibernating mammals shift metabolic fuel sources from carbohydrates to stored fat (Carey et al. 2003). Respiratory quotient estimates suggest lipids as the main fuel for hibernating black bears (Tøien et al. 2015). Consistently, elevated expression of genes involved in fatty acid mitochondrial oxidation was observed in the liver and bone marrow of hibernating black bears (Fedorov et al. 2011; Goropashnaya et al. 2021). However, we found transcriptional suppression of genes involved in fatty acid betta oxidation in muscle of hibernating black and brown bears that does not indicate elevated utilization of lipids as main fuel in muscle metabolism. This finding is in contrast to induction of lipid catabolism and fatty acid betta oxidation reported at transcriptional (Vermillion et al. 2015; Goropashnaya et al. 2020) and proteomic levels (Hindle et al., 2011) in muscle of hibernating 13-lined and Arctic ground squirrels. Thus, small mammalian hibernators and bears show considerable difference in transcriptional signature of muscle lipid metabolism during hibernation.

We found a coordinated transcriptional reduction of genes involved in complement and coagulation cascades in muscle of hibernating black bears. Similarly, transcriptional down regulation of innate and adaptive immunity genes was reported in bone marrow of black bears during hibernation (Goropashnaya et al. 2021). This finding implies suppression of innate immunity and coagulation that is supported by decrease in immune cell numbers (Sahdo et al. 2013; Graesli et al. 2015) and platelet aggregation (Fröbert et al. 2010) in blood of hibernating brown bears.

In conclusion, this study reveals coordinated transcriptional changes suggesting the active protein biosynthesis and suppression of autophagy dependent protein degradation through the mTORC1 mediated signaling as response to BCAA availability. These changes likely underlie an adaptation for maintaining muscle protein content through prolonged periods of immobility and fasting during winter hibernation. Transcriptional reduction of BCAA degradation implies preservation of BCAA content that supports protein biosynthesis through activating the mTORC1 signaling during fasting in hibernation. Down regulation of multiple genes involved in fuel metabolism is consistent with metabolic suppression and lower energy demand in hibernating bears. These changes in gene expression represent the transcriptional program common for hibernation of two species, black, and brown bears.

In line with the common functional genomics approach, these conclusions are based on genome-wide transcriptiome screening that makes testable predictions for follow up studies. The follow up studies need to include metabolomics estimates of BCAA content, phosphor-proteomic assessment of key downstream targets (*RPS6KB1*, *RPS6*) of the mTORC1 signaling involved in translation initiation and estimating protein biosynthesis rates in hibernating muscle by measuring incorporation of deuterium water (Miller et al. 2015).

## Acknowledgements

This work was supported by an Institutional Development Award (IDeA) from the National Institute of general Medical Sciences of the National Institutes of Health under grant number, [P20GM103395] and Center of Biomedical Research Excellence under grant number [P20GM130443]. The content is solely responsibility of the authors and does not necessarily represent the official view of the National Institutes of Health.

## Data Accessibility and Benefit-Sharing

Transcriptome sequencing data were archived on the NCBI Short Read Archive (BioProject XXX).

Benefits Generated: Benefits from this research accrue from the sharing of our data and results on public databases as described above.

## Author Contributions

VBF, AG and AVG designed research, VBF, AG, ØT, BMB, AVG conducted research, and data analysis. VBF and AVG wrote the paper with approval from all authors.

